# Δ^9^-tetrahydrocannabinol (THC) vapor exposure produces conditioned place preference in male and female rats

**DOI:** 10.1101/2021.12.20.473475

**Authors:** Catherine F. Moore, Catherine M. Davis, Cristina Sempio, Jost Klawitter, Uwe Christians, Elise M. Weerts

**Affiliations:** Division of Behavioral Biology, Department of Psychiatry and Behavioral Sciences, Johns Hopkins University School of Medicine; Department of Pharmacology and Molecular Therapeutics, Uniformed Services University of the Health Sciences, Bethesda, MD; iC42 Clinical Research and Development, Department of Anesthesiology, University of Colorado, Anschutz Medical Campus, Aurora, CO 80045

**Author notes:** Corresponding author at: Johns Hopkins Bayview Research Campus, Behavioral Biology Research Center, 5510 Nathan Shock Drive, Suite 3000, Baltimore, MD 21224, USA. E-mail address (C.F. Moore).

**Keywords:** THC, Vapor Exposure, Conditioned Place Preference, Reward, Cannabis

## Abstract

**Background:** The use of place conditioning procedures and drug vapor exposure models can increase our understanding of the rewarding and aversive effects of vaped cannabis products. Currently there are limited data on the conditioned rewarding effects of vaporized Δ^9^-tetrahydrocannabinol (THC), the primary psychoactive constituent of cannabis in rats, and no studies to date examining sex differences.

**Methods:** Male and female Sprague-Dawley rats (N=96; 12 per sex/group) underwent place conditioning sessions immediately after exposure to THC or vehicle (propylene glycol, PG) vapor. Locomotor activity was measured by beam breaks during conditioning sessions. THC vapor-conditioned rats received one of three THC vapor exposure amounts (low: 5 puffs of 100 mg/ml THC, medium: 5 puffs of 200 mg/ml THC, or high: 10 puffs of 200 mg/ml THC) and matched vehicle vapor (PG) exposure on alternate days for 16 daily sessions. A ‘no THC’ control group of vehicle-conditioned rats received only PG vapor exposure each day. After the 8th and 16th conditioning sessions, untreated rats were tested for conditioned place preference (CPP) or avoidance (CPA). Next, extinction tests and a THC vapor-primed reinstatement test were conducted.

**Results:** THC vapor produced CPP and locomotor effects in an exposure dependent manner, and some sex differences were observed. Low THC vapor exposure did not produce CPP in males or females. Medium THC vapor exposure produced CPP in males, but not females. High THC vapor exposure produced CPP in both males and females. Medium and high THC vapor exposure amounts produced hyperactivity in female rats, but not male rats. CPP was more resistant to extinction in females than males. THC vapor re-exposure (i.e., drug-prime) after extinction did not result in reinstatement of CPP for either sex.

**Conclusion:** This study demonstrates conditioned rewarding effects of THC vapor in both male and female rats and provides evidence for sex differences in amounts of THC vapor that produce CPP and in time to extinction. CPA was not observed at any of the THC vapor exposure amounts tested. These data provide a foundation for future exploration of the conditioned effects of cannabis constituents and extracts using vapor exposure models.

## Introduction

Vaping of cannabis and cannabinoids such as Δ^9^-tetrahydrocannabinol (THC, the primary psychoactive constituent of cannabis) is on the rise^1-4^. Recent estimated rates of vaping cannabis are reported to be between 20%-37% (past 30-days) and 60% (lifetime) in cannabis users in the US^5-7^. Vaping has also been promoted as a safer alternative to smoked cannabis ^8^. At the same time, a wide array of new products of cannabis constituents for vaping are now available, with little data available on potential abuse liability.

Place conditioning procedures can reveal the conditioned rewarding or aversive effects of an administered substance^9^. In this procedure, drug is administered followed with pairing in a distinct environment with various contextual clues. On alternate conditioning sessions, vehicle administration (no drug) is paired with a separate, distinct environment. On a drug- and vehicle-free test day, animals are allowed unrestricted access to both environments to assess conditioned place preference (CPP) or aversion (CPA). Increases in time spent on the drug-paired side (i.e., preference) indicates conditioned rewarding effects of the drug, and decreases in time spent on the drug-paired side indicates conditioned aversive effects of the drug.

Following the acquisition of CPP, repeated, drug-free extinction tests can be conducted to assess the persistence of preference to the conditioned chamber^10^. To assess relapse-like behavior, reinstatement test can be conducted following exposure to a stress-, cue-, or drug-induced priming^11^.

To date, the majority of research examining the conditioned rewarding effects of cannabis and cannabinoids have primarily used the intraperitoneal (IP) injection route of administration ^9^. When injected IP, low THC doses (0.075-4 mg/kg) can produce preference, while higher THC doses (5-20 mg/kg) produce aversion for reviews, see^12, 13^. To date, one study has tested for CPP with THC vapor exposure in male rats; one out of the six conditions tested produced a preference, while the other five vapor conditions produced neither CPP nor CPA^14^.

Depending on route of administration, THC can produce markedly different physiological and behavioral effects, due to distinctive pharmacokinetics and metabolic pathways^15, 16^. For example, oral and IP administered THC undergoes first pass metabolism in the liver, whereas vaped or intravenously administered THC does not. Previously, we evaluated the antinociceptive, hypothermic, and appetitive effects of a range of THC vapor exposure conditions in comparison to IP administered THC in two strains of male and female rats^17^. We demonstrated effects of THC vapor were orderly and exposure dependent, and that THC vapor produced plasma concentrations of THC that were comparable to those observed in humans.

Out of 5 conditions tested, lower amounts of THC vapor exposure increased appetitive effects in a test of progressive ratio responding for food pellet rewards, while the higher amounts of THC vapor exposure resulted in hypothermia, antinociception, and sedative effects^17^. The current study sought to evaluate whether the THC vapor exposure conditions equivalent to those that produced appetitive effects in our prior study (previously called THC exposure conditions 1, 2, and 3 – here designated low, medium, and high) would produce CPP. The current study design included both male and female rats and an ‘exposure-effect’ curve with multiple THC exposure groups to assess CPP. Second, we evaluated extinction and drug-primed reinstatement of CPP. In a separate group of rats, we analyzed levels of THC and key metabolites, 11-hydroxy-THC (11-OH-THC) and 11-Nor-9-carboxy-THC (THC-COOH) after THC exposure conditions, to confirm relevant levels of drug exposure.

## Materials and Methods

### Subjects

Adult male and female Sprague Dawley rats (N=96, 12 per sex/group) (Charles River, Wilmington, MA), 8 weeks old at the start of experiments, were single housed in wire-topped, plastic cages (27 × 48 × 20 cm) with standard enrichment. The vivarium was on a 12 hr reverse light cycle (lights off at 8:00 a.m.) and was humidity and temperature controlled. Diet was a corn-based chow (Teklad Diet 2018; Harlan, Indianapolis, IN); rats had ad libitum access to food and water except during test procedures. All procedures used in this study were approved by the Johns Hopkins Institutional Animal Care and Use Committee. The facilities adhered to the National Institutes of Health *Guide for the Care and Use of Laboratory Animals* and were AAALAC-approved.

### Drugs

THC stock solutions (100 and 200 mg/ml in 95% ethanol), confirmed ≥95% purity, were provided by the U.S. National Institute on Drug Abuse Drug Supply Program. Ethanol from the THC stock was evaporated using nitrogen and then THC was mixed in 100% propylene glycol to yield a 100 or 200 mg/ml THC solution for vaporization.

### Place conditioning and testing

The conditioning apparatus used contained 3-chambers, each with distinct environments and automated recording of activity via infrared beam breaks (San Diego Instruments, San Diego, CA). The left chamber had white walls and a textured plastic floor, the middle chamber had clear walls and a stainless steel, grid rod floor, and the right chamber had black walls and a smooth plastic floor. To evaluate the potential rewarding effects of THC vapor exposure, testing included 3 phases, pre-test, conditioning, and post-tests. In the pre-test, rats were placed in the center chamber and allowed to freely explore the 3-chamber apparatus for 15 min to assess normal side bias prior to conditioning. Overall, no significant group bias was present (Supplemental Fig S1; Supplemental Table S1). Rats were then allocated into four test groups and balanced based on individual pretest CPP score (detailed in the data analysis section below). For the conditioning phase, rats received either 100% propylene glycol (‘no THC’ condition), or one of three different THC vapor exposure amounts (low, medium, high) as detailed in Table 1, using a commercial vapor chamber system (La Jolla Alcohol Research Institute, La Jolla, CA) fitted with an electronic vape device (Smok Baby Beast Brother TFV8 Sub-Ohm Tank with the V8 X-Baby M2 0.25-Ω coil; SMOKTech, Shenzhen, China) as described in detail previously^17^. Rats were passively exposed to either the assigned THC vapor exposure amount or a matched vehicle vapor condition on alternating days (THC, VEH, THC etc.; see Fig 1) and confined to the left or right chamber for 30 min (THC vapor = non-preferred chamber; vehicle vapor = preferred chamber). Vehicle-conditioned rats received vehicle vapor exposure each day. Rats underwent two sets of 8 daily conditioning sessions (16 days total).

**Table 1.** Amounts of vapor exposure for each group. Each puff was delivered at equally set intervals (e.g., 2.5-5 min apart) across the 30-min session. Estimated total THC drug amounts (mg) delivered per exposure session were calculated based on volume delivered (determined by tank weights before and after sessions) and THC concentration in e-liquid; Note: air flow is continuous and vaporized drug is pulled out into the exhaust system.

| Vapor Group | N (50% female) | Vehicle Conditioning Days |  |  | THC Conditioning Days |  |  |  |
| --- | --- | --- | --- | --- | --- | --- | --- | --- |
|  |  | VEH | Puffs (# x length) | Exposure Duration (min) | THC Concentration (mg/ml) | Puffs | Exposure Duration (min) | Calculated THC dose (mg) |
| No THC | 24 | 100% PG | 5 x 6s | 30 | 0 (100% PG) | 5 x 6s | 30 | 0 |
| Low THC | 24 | 100% PG | 5 x 6s | 30 | 100 | 5 x 6s | 30 | 7 |
| Medium THC | 24 | 100% PG | 5 x 6s | 30 | 200 | 5 x 6s | 30 | 10 |
| High THC | 24 | 100% PG | 10 x 6s | 30 | 200 | 10 x 6s | 30 | 17 |

**Figure 1.**
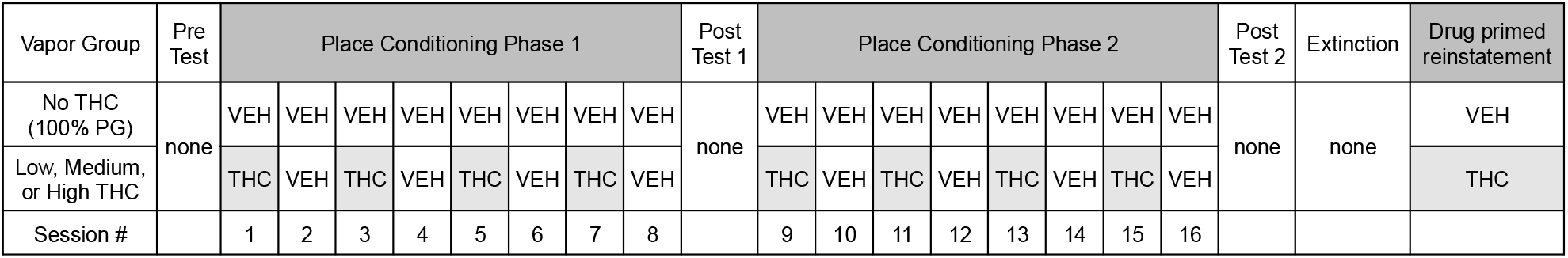
Conditioned place preference testing timeline. Rats were conditioned across 16 days of vapor exposure with a preference test on day 0, between conditioning days 8-9 and after conditioning day 16.

Using the same procedures as the pre-test, described above, rats were tested for expression of CPP in a drug-free state after the 8^th^ conditioning session (post-test 1) and the 16^th^ conditioning session (post-test 2). As detailed in the analysis section, the criteria for CPP for the THC vapor-paired chamber, were defined *a priori*. In the event of demonstration of CPP after post-test 2, daily drug-free tests continued for that group until extinction criteria were met (see data analysis section). Vehicle vapor control groups were run for an equal number of extinction days as THC groups for comparison. Following extinction, each rat was exposed to a single vapor session using the same exposure parameters they received during the conditioning phases to determine if THC vapor-primed reinstatement of CPP; prior to a 15-min preference test (see Fig 1 for a timeline of test procedures).

### Analysis of THC and key metabolites after THC vapor exposure

Separate groups of rats (N=4/sex) were exposed to selected THC vapor conditions used in the CPP study (medium and high); rats were euthanized via isoflurane overdose and blood was collected into EDTA tubes approximately 10-min after THC vapor exposure. Blood samples were placed on wet ice for 30 min prior to centrifugation at 3000x*g* for 10 min. Plasma supernatant was transferred to low protein binding microcentrifuge tubes and stored at −80°C until shipment on dry ice to iC42 Clinical Research and Development (Aurora, CO) where THC and its key metabolites 11-Hydroxy-Δ^9^-tetrahydrocannabinol (11-OH-THC) and 11-Nor-9-carboxy-Δ^9^-tetrahydrocannabinol (THC-COOH) were quantified using an established validated high-performance liquid chromatography-tandem mass spectrometry (LC-MS/MS) assay.

Details of the assay and validation results are as previously described^18^. The results of blank, zero, calibrators and quality control samples included in the study sample batch met all predefined acceptance criteria: the calibration range for THC and THC-COOH was 0.78-400 ng/mL and that for 11-OH-THC 1.56-400 ng/ml. There was no carry over and no matrix interferences. Accuracy in the study sample batch was within the ±15% acceptance criterion and imprecision was <15%.

### Data Analysis

A CPP score was calculated (time in THC vapor paired chamber – time in unpaired chamber). Criteria for determination of a place preference and extinction were defined *a priori* as a positive change in preference score (CPP Score Post-Test – CPP Score Pre-Test) > 0, determined by one-sample t-test. Extinction criteria were defined as two consecutive days of no preference (change in preference = 0, determined by one-sample t-test). Partial eta squared (η_p_^2^) was used to report effect sizes (>0.14 is considered a large effect size).

To assess whether locomotor activity was altered after acute and intermittent THC exposure vs. vehicle vapor exposure, we analyzed distance traveled during conditioning sessions on day 1, day 7, and day 15. For locomotor activity analysis, effects of exposure group and sex (between subjects effects) and day (repeated within subjects effect) were analyzed with a three-way ANOVA. In the event of a three-way interaction, two-way ANOVAs were run in each level of the third factor and post hoc Holm–Sidak tests for multiple comparisons were run to interpret significant interactions.

Two-way ANOVAs were used to assess sex and group effects on plasma levels of THC and 11-OH-THC. Statistics were performed in Statistica 11 (StatSoft, Tulsa, OK) and GraphPad Prism 9 (GraphPad Software, San Diego, CA) with p ≤ 0.05 for significance. The data were Greenhouse-Geisser corrected where Mauchly’s Sphericity tests were significant.

## Results

### Development of a preference for THC vapor-paired chamber

After each post-test, preference was determined for each group using the *a priori* criteria described above. In males, none of the exposure amounts resulted in a change in CPP score tested after 8 days of conditioning (i.e. four THC and four vehicle vapor exposures); however after 16 days of conditioning, CPP was demonstrated in males in the medium THC vapor exposure group [t(11)=2.940, p=0.0135, η_p_^2^= 0.44] and the high THC vapor exposure group [t(11)=4.549, p=0.0008, η_p_^2^= 0.65)](Fig 2). In females, 8 days of conditioning to high THC vapor was sufficient to produce place preference for the THC vapor-paired chamber (t(11) = 2.959, p=0.0130, η_p_^2^= 0.44). After continued conditioning, CPP was still evident in this group at Post-Test 2 (t(11) = 2.821, p=0.0166, η_p_^2^= 0.42). CPP was not observed after 8 or 16 days of conditioning in females in the low and medium THC vapor exposure groups (Fig 2A-B).

**Figure 2.**
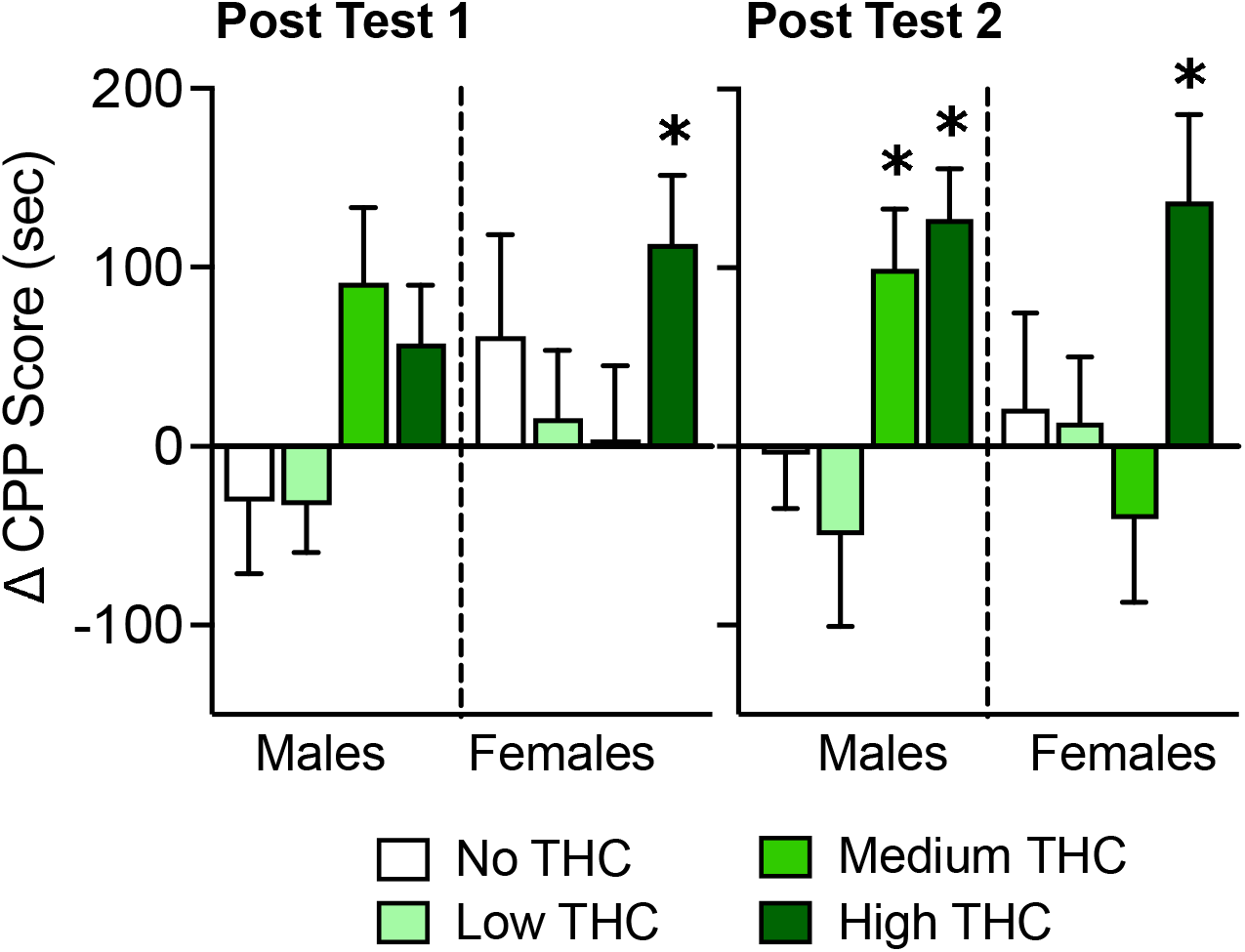
Change in preference score across successive place preference tests. Data shown are group means (±SEM) for post-tests 1 and 2, conducted after each conditioning days 1-8 and 9-16, respectively. An asterisk * represents a place preference (CPP Score Post-Test – CPP Score Pre-Test > 0; p>0.05, determined by a one-sample t-test).

Male and female rats in the vehicle vapor conditioning group did not show increases or decreases in the time spent in the initially unpreferred (i.e., drug-paired for THC groups) chamber (Supplemental Fig S1).

### Extinction and reinstatement tests

Males in the medium THC vapor group showed a transient increase in preference on day 1 of extinction testing (t(11) =3.94, p<0.01; η_p_^2^= 0.59) that decreased with additional sessions (p’s>0.05 on extinction day 2 and 3) (Fig 3). Males in the high THC vapor group showed immediate decrease in CPP (p’s > 0.05s) on extinction day 1 and 2. Females in the high THC vapor group showed an increase in preference for the THC vapor-paired chamber on days 1 and 2 of extinction testing, which decreased on the 3^rd^ and 4^th^ extinction tests (Extinction day 1: t(11) = 3.36, p<0.01, η_p_^2^= 0.51; Extinction day 2: t(11) = 3.11, p<0.01, η_p_^2^= 0.56)(Fig 2C).

**Figure 3.**
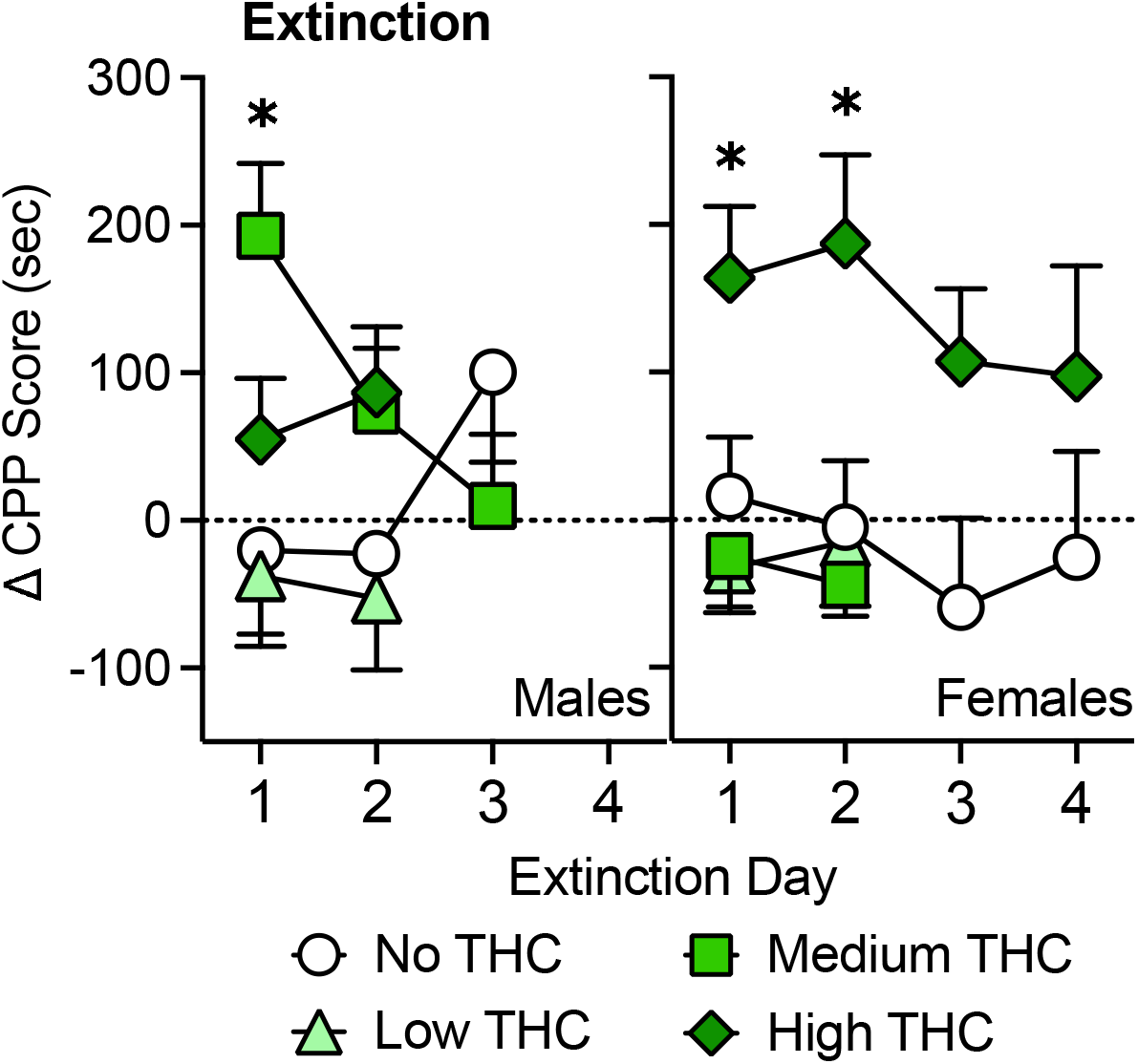
Change in preference score across successive extinction tests. Data shown are group means (±SEM). An asterisk * represents a place preference (CPP Score Post-Test – CPP Score Pre-Test > 0; p>0.05, determined by a one-sample t-test).

However, upon closer analysis of the data following the end of the experimental protocol, one female in the high THC group was identified as a consistent outlier (>2 standard deviations *below* group mean on Post-Test 2 and Extinction days 1-4). If this female is removed, the mean change in CPP score for the remaining females on the last day tested is 152.7 ±54.09 (t(10) =2.82, p=0.02), which suggests that resistance to extinction of CPP in females was even more prolonged.

In groups that displayed CPP, the day following the final extinction day, animals were reexposed to their CPP exposure amount of THC vapor prior to a preference test. This drug-prime did not result in reinstatement of CPP to the THC vapor-paired chamber in any group (p’s>0.05; group means (±SEM) were Males: no THC = -14.11 (±52.3), medium THC = -7.30 (±80.7), high THC= 79.65 (±69.8); Females: no THC = -95.02 (±62.5), high THC = 4.13 (±43.9) (data not shown).

### Locomotor effects of acute and repeated THC vapor exposure

In a three-way ANOVA of activity across the conditioning days (1, 7, 15), with sex and group as between subjects effects, Mauchly’s test of sphericity was significant, χ^2^(2) = 11.06, p = 0.004, and Greenhouse-Geisser correction is presented. We observed a three-way interaction of Day x Sex x Group (F (5.36,157.24) = 3.28, p<0.01; Fig 4).

**Figure 4.**
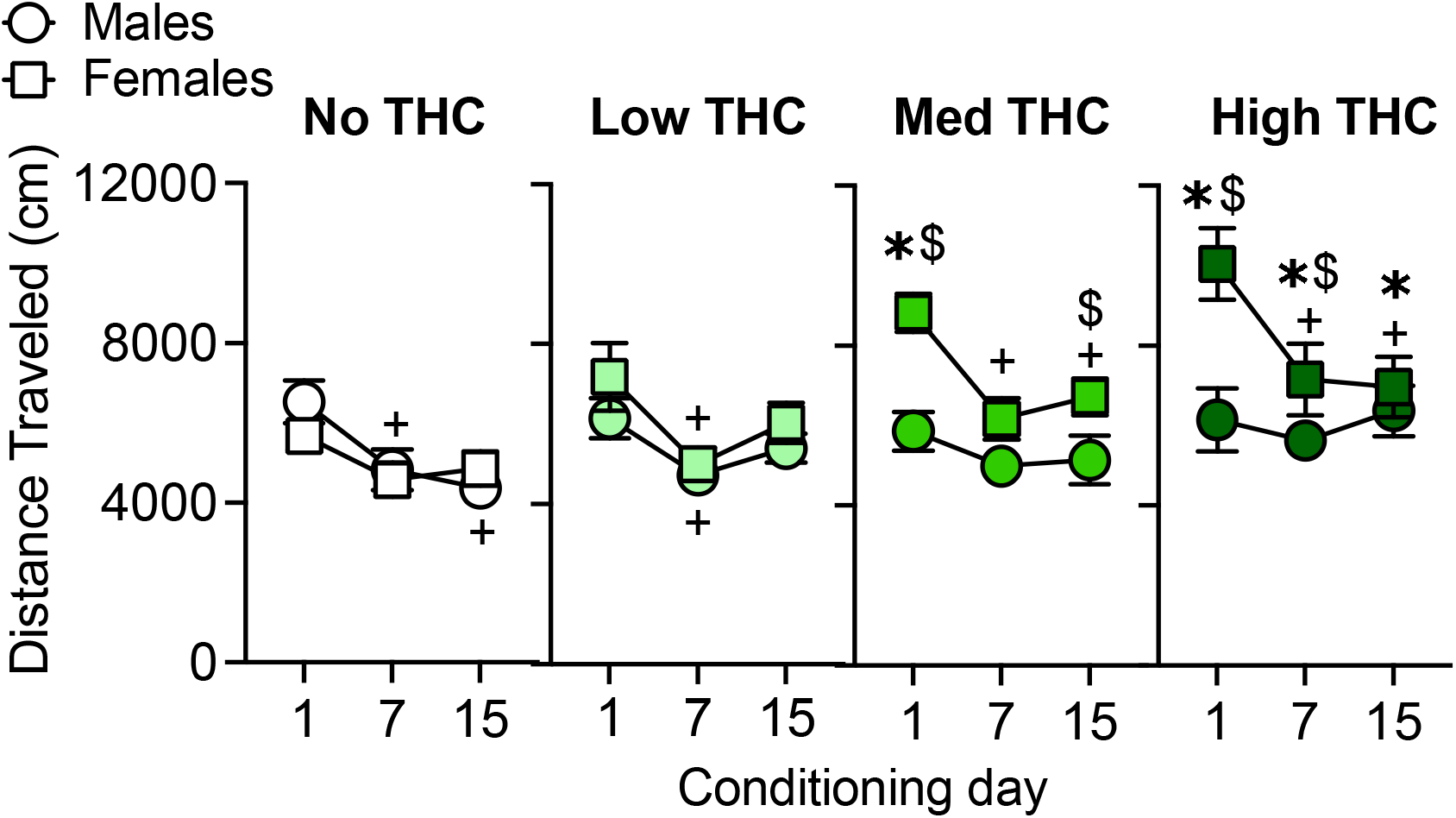
Distance traveled on conditioning days 1, 7, and 15. Asterisks (*) denote significant post-hoc differences between a THC group compared to the vehicle (‘no THC’) vapor control group on that day; dollar signs ($) denote significant post-hoc sex differences on that day. Plus signs (+) denote differences from Day 1 for that group (p’s<0.05).

Post-hoc tests indicate that females in the medium and high THC vapor groups had higher locomotor activity compared to males of the same THC vapor group on the first day of vapor exposure (medium THC group, p<0.01, Cohen’s *d* = 1.78; high THC group, p<0.001, Cohen’s *d* = 1.33). On Day 7, females in the high THC vapor group continued to have higher locomotor activity post-THC vapor exposure compared with males of the same drug condition (p<0.01, Cohen’s *d* = 0.62). On Day 15, the medium THC vapor females had higher locomotor activity post-THC vapor exposure compared with males of the same drug condition (p<0.05, Cohen’s *d* = 0.84).

In females, but not males, THC exposure dose-dependently increased locomotor activity. Post-hoc tests indicate that locomotor activity of females in medium and high THC vapor groups were higher on Day 1 of THC exposure (Day 1: medium group, p<0.01, Cohen’s *d* = 2.18; high THC group, p<0.001, Cohen’s *d* = 1.85). On Days 7 and 15, the high THC group was still increased compared with vehicle (Day 7, p<0.01, Cohen’s *d* = 1.06; Day 15, p<0.05, Cohen’s *d* = 0.99). In males, there were no significant differences in locomotor activity between Vehicle and THC vapor groups; though on Day 15, the high THC group tended to have higher locomotor activity than the vehicle vapor group (p=0.056; Cohen’s *d* = 1.09).

Post-hoc tests evaluating changes in locomotor response to THC across the conditioning days (i.e., tolerance or sensitization) demonstrated that females in the medium and high THC vapor groups had higher activity on Day 1 compared to Day 7 (medium: p<0.001, Cohen’s *d* = 1.55; high: p<0.001, Cohen’s *d* = 0.93) and Day 15 (medium: p<0.01, Cohen’s *d* = 1.30; high: p<0.001, Cohen’s *d* = 1.07). Males in the vehicle vapor group and males and females in the low THC vapor group also showed lower activity on Day 7 compared with Day 1 (Males Vehicle vapor: p<0.01, Cohen’s d = 0.94; Males low THC vapor: p<0.05, Cohen’s *d* = 0.94; females low THC vapor: p<0.001, Cohen’s *d* = 0.94). Males in the vehicle vapor group also had lower activity on Day 15 compared with Day 1 (p<0.01, Cohen’s *d* = 1.30). Therefore, locomotor activity decreased (i.e., habituated) across days in some groups that did not show initial (Day 1) increases in activity after vapor exposure.

### Blood concentrations of THC and metabolites

Analysis of blood samples from rats exposed to medium and high THC vapor exposure conditions revealed exposure dependent increases in levels of THC and key metabolites, 11-OH-THC and THC-COOH (Figure 5).

**Figure 5.**
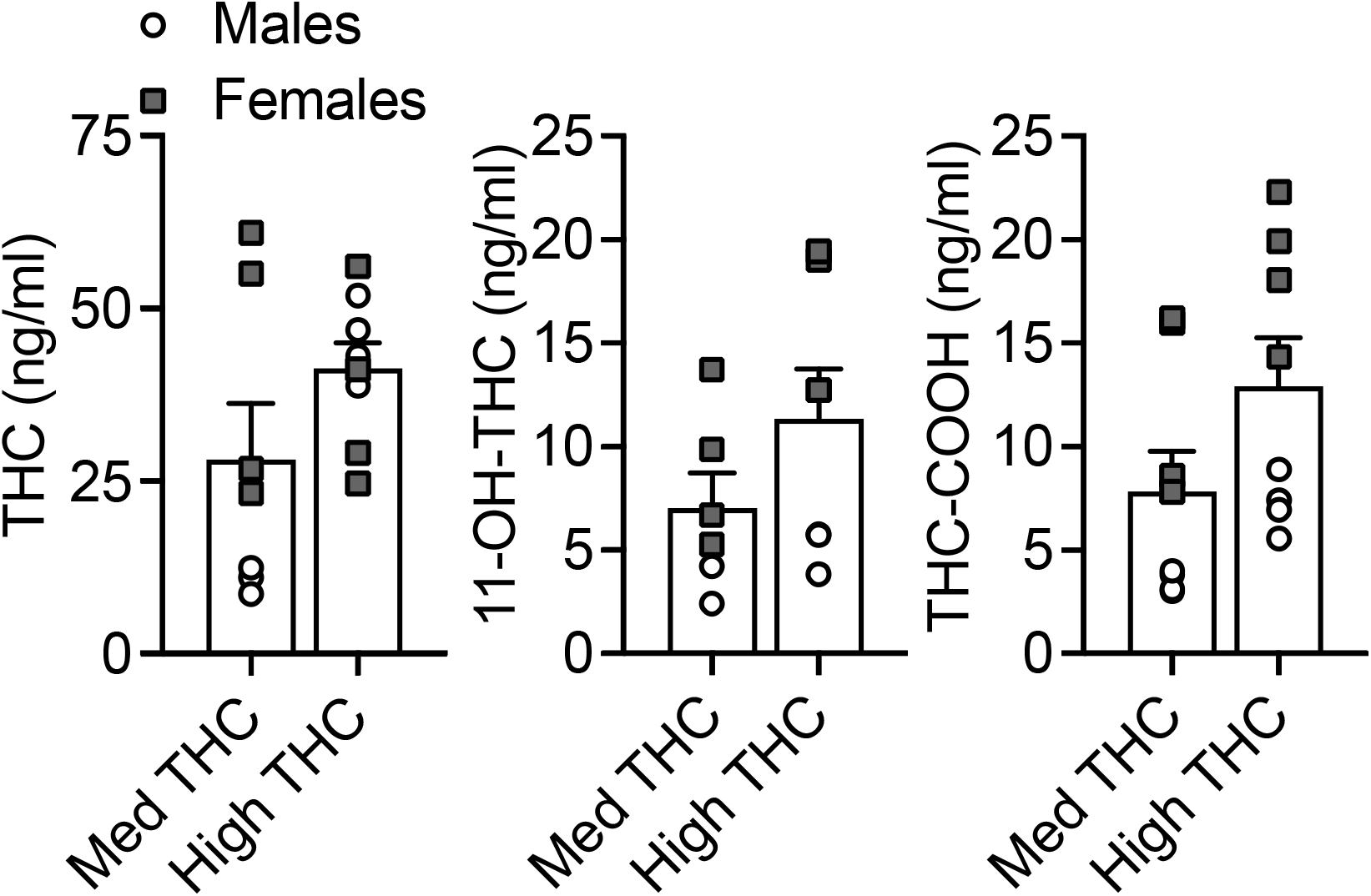
Levels of THC, 11-OH-THC, and THC-COOH in plasma of rats after acute exposure to medium and high THC vapor.

## Discussion

In this study, we observed THC vapor-induced CPP in male and female rats although there were sex differences based on exposure amounts and duration to extinction. These data add to prior research examining conditioned effects of THC administered via IP injection. While many prior studies using different THC doses administered IP have reported CPP, CPA, or null effects, a recent meta-analysis determined that CPP is most often observed after lower doses (median dose 1 mg/kg: range 0.075-4 mg/kg) whereas higher doses produce CPA (median dose 5 mg/kg; range: 1-20 mg/kg)^13^. Dose-dependent effects of IP injected THC on anxiety-like behaviors mirror what has been seen with CPP/CPA; low doses produce anxiolytic effects while high doses produce anxiogenic effects^19, 20^. Neurobiologically, low doses of injected THC have been shown to increase extracellular dopamine in reward-related brain regions including the nucleus accumbens shell^21^ and striatum^22^; whereas higher doses of THC increase corticosterone via centrally mediated mechanisms^23-25^. Therefore, the dose-dependency observed in CPP experiments with injected THC tracks with the dose-dependent effects of THC observed in related behavioral and neurobiological outcomes. In the current study, we chose exposure amounts based on our previous study comparing IP injected THC with vaporized THC conditions. In these studies, behavioral outcomes of THC exposure conditions 1, 2, and 3 (corresponding to low-medium-high in the present study) were comparable to _≲_1-3 mg/kg THC injected IP.

In rats exposed to medium and high THC vapor, mean plasma levels of THC were 28.3 and 41.4 ng/ml, respectively. In comparison, two human laboratory studies using vaporized cannabis resulted in a C_max_ of 14 ng/ml THC (25 mg dose)^26^ and 47 ng/ml THC (50 mg dose)^27^. Our ranges of THC levels in plasma are comparable to what has been shown after IP injection of low doses of THC 0.5-3 mg/kg (∼25-75 ng/ml)^28-30^, after passive THC vapor exposure (35-73 ng/ml)^17, 28, 31^, and in rats self-administering THC vapor puffs (∼10-40 ng/ml)^32, 33^. Blood levels of THC following vapor exposure peak rapidly, within 5-10 minutes after exposure^28, 31^. Blood levels of the key THC metabolites 11-OH-THC and THC-COOH were relatively low 7-11 and 8-13 ng/ml, respectively, mirroring what has been shown previously with vapor exposure models^28^. As 11-OH-THC and THC-COOH are bioactive metabolites and have activity at the cannabinoid type-1 receptors, their pharmacokinetic profile with respect to vapor exposure is an important factor in THC rewarding effects.

To date, only one prior study has examined conditioned effects of THC vapor, and the study was limited to one sex (male rats). In this study, one THC vapor exposure amount (10 mg in 16L air, delivered over 10-min) prior to conditioning sessions produced a modest preference for the THC-vapor paired side^14^. Other exposure conditions tested did not produce CPP or CPA. Specifically, rats exposed to the same dose (10 mg in 16L air) but delivered over a longer period (20-min) did not show a preference for the THC-paired side^14^. Additionally, a group receiving the same THC dose over the same duration (10 mg over 10-min), but administered to two rats simultaneously and in a reduced volume (8L air), also did not produce preference, potentially due to less total dose exposure (10 mg/2 rats vs. 10 mg/1 rat)^14^. Similar to our study, no THC vapor exposure amount tested produced aversion. It is unclear if this is related to route of administration (e.g., vapor exposure vs. injected THC), or vapor exposure conditions (e.g., THC ‘dose’ or duration of exposure tested) to produce CPA. We intend on addressing this question in future studies.

We are aware of one evaluation of sex differences in THC CPP, in which THC was administered via IP injection^34^; a place aversion to THC (0.75 mg/kg) was observed but there was no difference between in expression of CPA in males and females. In the present study, we observed CPP to THC vapor in both males and females, but female rats required a greater amount of THC vapor to produce CPP compared with males. This finding was somewhat surprising, as the literature suggests that females are more sensitive to cannabinoid effects^35^, including the reinforcing^36, 37^ and discriminative effects^38^ of synthetic cannabinoid receptor type-1 agonists. Indeed, in our previous study, females showed increased appetitive responses after a lower amount of THC vapor exposure compared with males^17^. Consistent with an increased sensitivity to THC in females, locomotor stimulating effects of THC vapor were evident in females, but not males in the current study. In females, although a higher amount of THC was required for development of a CPP, this was evident after only 8 conditioning days. In contrast, in males, the medium and high THC condition produced an increase in CPP that was below the significance threshold after 8 days, but further increased after 8 additional conditioning days (16 total). This would indicate that females acquire conditioned rewarding effects of THC somewhat faster than males, despite requiring higher exposure amounts. It may be that initial exposures to vaporized THC are aversive (particularly in males), requiring additional conditioning days to overcome this early experience. This theory is based on a prior study investigating CPP to injected THC, where a protocol implementing an initial priming injection of THC shifted outcomes from no preference to a preference (1 mg/kg) or from aversion to no preference (5 mg/kg) in primed vs. naive male mice^39^. This shift in CPP was assumed to result from a disassociation of the initial aversive drug effects with the later drug-paired CPP context^13^. To date no preclinical studies have observed any aversive qualities of acute THC vapor; a study using intra-cranial self-stimulation (ICSS) found that cannabis smoke exposure caused a mild reduction in the reward threshold (i.e. dysphoric effects), though only after repeated exposures (<4 days)^40^. Clinical evidence to date would suggest that females are more sensitive to the anxiogenic effects of acute cannabinoids compared with males^41^; to date, there are very few studies of sex differences in acute THC effects on anxiety-like or other aversive behaviors^42^.

Following cessation of THC vapor conditioning, we observed extinction to the conditioned rewarding effects in all groups within 4 days. Females demonstrated a greater resistance to extinction compared with males, who extinguished CPP to THC vapor more rapidly. We did not observe drug-primed reinstatement following extinction of CPP in either males or females. It is possible that a higher amount of THC vapor may be needed to induce reinstatement, due to increased tolerance resulting from chronic THC vapor administration. Alternatively, lower doses may be advantageous for observing reinstatement as is the case with certain drugs, including ethanol^43, 44^. This was the first study that assessed drug primed reinstatement with THC and was limited to one exposure condition. Future studies should explore different exposure amounts, as well as cue or stress-primed reinstatement of CPP.

Our study used a biased conditioning procedure (i.e., biased stimulus assignment procedure^45^); the THC vapor was paired with the initially non-preferred chamber in an attempt to ‘switch’ the preference. Thus, it is important to consider that with repeated exposures, an initially non-preferred chamber may lose its aversive qualities resulting in a shift in bias unrelated to drug effects. To control for this, we used a vehicle only control group, which received vehicle vapor (100% PG, no THC) prior to placement in either chamber, with equal numbers of conditioning sessions as THC vapor groups and run concurrently. We saw no change in the time spent in the initially non-preferred chamber in the vehicle vapor groups (Supplemental Figure S1B). There is a chance that THC vapor exposure simply reduced the aversive qualities of the initially non-preferred (now drug-paired) chamber, rather than produced a true preference. This cannot be completely ruled out. However, there is no evidence that our CPP apparatus is biased^45, 46^; group means show equal time spent in the left (black) and right (white) chambers (Supplemental Figure S1A). Overall, the preference produced by medium and high THC vapor exposure in our study would be considered moderate when compared to effects observed for some drugs like cocaine^47^, but is greater when compared to vehicle vapor groups.

## Conclusions

This study builds on prior research by demonstrating CPP after exposure to multiple amounts of THC vapor and durations of conditioning in both male and female rats. This is the first study to examine sex differences in the amount of THC exposure and number of conditioning sessions needed to produce CPP, as well time to extinction of CPP. Further, we saw a striking sex difference in locomotor response to THC vapor, in which in only females showed hyperlocomotion on THC vapor conditioning days. Thus, these data also add to the growing literature showing sex-related differences in sensitivity to the pharmacological effects of THC vapor.

## Abbreviations

11-OH-THC: 11-Hydroxy-Δ^9^-tetrahydrocannabinol
ANOVA: analysis of variance
CPP: Conditioned Place Preference
CPA: Conditioned Place Aversion
LC-MS/MS: liquid chromatography- tandem mass spectrometry
MED: Medium
PG: Propylene Glycol
SEM: standard error of the mean
THC: Δ^9^-tetrahydrocannabinol
THC-COOH: 11-Nor-9-carboxy-Δ^9^-tetrahydrocannabinol
VEH: Vehicle

## Acknowledgements

The authors wish to thank Maury Cole and La Jolla Alcohol Research Inc. for development of custom vapor chamber systems and technical assistance. The authors also would like to thank the NIDA Drug Supply Program for providing THC.

## Authorship contribution statement

### Catherine Moore

Investigation, Formal analysis, Writing - Original Draft; **Elise Weerts:** Conceptualization, Funding acquisition, Project administration, Writing - Review & Editing; **Catherine Davis**: Conceptualization, Writing - Review & Editing; **Cristina Sempio, Jost Klawitter, and Uwe Christians**: Investigation, Methodology, Validation, Writing - Review & Editing.

## References

1. Hasin DS, Borodovsky J, Shmulewitz D, et al. Use of highly-potent cannabis concentrate products: More common in U.S. states with recreational or medical cannabis laws. Drug and Alcohol Dependence 2021; 229(Pt B): 109159. doi:10.1016/j.drugalcdep.2021.109159.

2. Knapp AA, Lee DC, Borodovsky JT, et al. Emerging Trends in Cannabis Administration Among Adolescent Cannabis Users. Journal of Adolescent Health 2019; 64(4): 487–493. doi:10.1016/j.jadohealth.2018.07.012.

3. Morean ME, Kong G, Camenga DR, et al. High School Students’ Use of Electronic Cigarettes to Vaporize Cannabis. Pediatrics 2015; 136(4): 611–6. doi:10.1542/peds.2015-1727.

4. Varlet V, Concha-Lozano N, Berthet A, et al. Drug vaping applied to cannabis: Is “Cannavaping” a therapeutic alternative to marijuana? Sci Rep 2016; 6: 25599. doi:10.1038/srep25599.

5. Lee DC, Crosier BS, Borodovsky JT, et al. Online survey characterizing vaporizer use among cannabis users. Drug and Alcohol Dependence 2016; 159: 227–33. doi:10.1016/j.drugalcdep.2015.12.020.

6. Schauer GL, Njai R, and Grant-Lenzy AM. Modes of marijuana use - smoking, vaping, eating, and dabbing: Results from the 2016 BRFSS in 12 States. Drug and Alcohol Dependence 2020; 209: 107900. doi:10.1016/j.drugalcdep.2020.107900.

7. Goodman S, Wadsworth E, Leos-Toro C, et al. Prevalence and forms of cannabis use in legal vs. illegal recreational cannabis markets. International Journal of Drug Policy 2020; 76: 102658. doi:10.1016/j.drugpo.2019.102658.

8. Budney AJ, Sargent JD, and Lee DC. Vaping cannabis (marijuana): parallel concerns to e-cigs? Addiction 2015; 110(11): 1699–704. doi:10.1111/add.13036.

9. Murray JE and Bevins RA. Cannabinoid conditioned reward and aversion: behavioral and neural processes. ACS Chem Neurosci 2010; 1(4): 265–278. doi:10.1021/cn100005p.

10. Aguilar MA, Rodriguez-Arias M, and Minarro J. Neurobiological mechanisms of the reinstatement of drug-conditioned place preference. Brain Research Reviews 2009; 59(2): 253–77. doi:10.1016/j.brainresrev.2008.08.002.

11. Shaham Y, Shalev U, Lu L, et al. The reinstatement model of drug relapse: history, methodology and major findings. Psychopharmacology (Berl) 2003; 168(1-2): 3–20. doi:10.1007/s00213-002-1224-x.

12. Panagis G, Vlachou S, and Nomikos GG. Behavioral pharmacology of cannabinoids with a focus on preclinical models for studying reinforcing and dependence-producing properties. Curr Drug Abuse Rev 2008; 1(3): 350–74. doi:10.2174/1874473710801030350.

13. Kubilius RA, Kaplick PM, and Wotjak CT. Highway to hell or magic smoke? The dose-dependence of Delta(9)-THC in place conditioning paradigms. Learn Mem 2018; 25(9): 446–454. doi:10.1101/lm.046870.117.

14. Manwell LA, Charchoglyan A, Brewer D, et al. A vapourized Delta(9)- tetrahydrocannabinol (Delta(9)-THC) delivery system part I: development and validation of a pulmonary cannabinoid route of exposure for experimental pharmacology studies in rodents. Journal of Pharmacological and Toxicological Methods 2014; 70(1): 120–7. doi:10.1016/j.vascn.2014.06.006.

15. Hlozek T, Uttl L, Kaderabek L, et al. Pharmacokinetic and behavioural profile of THC, CBD, and THC+CBD combination after pulmonary, oral, and subcutaneous administration in rats and confirmation of conversion in vivo of CBD to THC. Eur Neuropsychopharmacol 2017; 27(12): 1223–1237. doi:10.1016/j.euroneuro.2017.10.037.

16. Spindle TR, Cone EJ, Goffi E, et al. Pharmacodynamic effects of vaporized and oral cannabidiol (CBD) and vaporized CBD-dominant cannabis in infrequent cannabis users. Drug and Alcohol Dependence 2020; 211: 107937. doi:10.1016/j.drugalcdep.2020.107937.

17. Moore CF, Davis CM, Harvey EL, et al. Appetitive, antinociceptive, and hypothermic effects of vaped and injected Delta-9-tetrahydrocannabinol (THC) in rats: exposure and dose-effect comparisons by strain and sex. Pharmacology Biochemistry and Behavior 2021; 202: 173116. doi:10.1016/j.pbb.2021.173116.

18. Sempio C, Almaraz-Quinones N, Jackson M, et al. Simultaneous Quantification of 17 Cannabinoids by LC-MS-MS in Human Plasma. Journal of Analytical Toxicology 2022; 46(4): 383–392. doi:10.1093/jat/bkab030.

19. Rubino T, Sala M, Vigano D, et al. Cellular mechanisms underlying the anxiolytic effect of low doses of peripheral Delta9-tetrahydrocannabinol in rats. Neuropsychopharmacology 2007; 32(9): 2036–45. doi:10.1038/sj.npp.1301330.

20. Valjent E, Mitchell JM, Besson MJ, et al. Behavioural and biochemical evidence for interactions between Delta 9-tetrahydrocannabinol and nicotine. British Journal of Pharmacology 2002; 135(2): 564–78. doi:10.1038/sj.bjp.0704479.

21. Tanda G, Pontieri FE, and Di Chiara G. Cannabinoid and heroin activation of mesolimbic dopamine transmission by a common mu1 opioid receptor mechanism. Science 1997; 276(5321): 2048–50. doi:10.1126/science.276.5321.2048.

22. Malone DT and Taylor DA. Modulation by fluoxetine of striatal dopamine release following Delta9-tetrahydrocannabinol: a microdialysis study in conscious rats. British Journal of Pharmacology 1999; 128(1): 21–6. doi:10.1038/sj.bjp.0702753.

23. Kubena RK, Perhach JL, Jr., and Barry H, 3rd. Corticosterone elevation mediated centrally by delta 1-tetrahydrocannabinol in rats. European Journal of Pharmacology 1971; 14(1): 89–92. doi:10.1016/0014-2999(71)90128-2.

24. Kokka N and Garcia JF. Effects of delta 9-THC on growth hormone and ACTH secretion in rats. Life Sciences 1974; 15(2): 329–38. doi:10.1016/0024-3205(74)90223-9.

25. DeVuono MV, La Caprara O, Petrie GN, et al. Cannabidiol Interferes with Establishment of Delta(9)-Tetrahydrocannabinol-Induced Nausea Through a 5-HT1A Mechanism. Cannabis Cannabinoid Res 2020. doi:10.1089/can.2020.0083.

26. Spindle TR, Cone EJ, Schlienz NJ, et al. Acute Pharmacokinetic Profile of Smoked and Vaporized Cannabis in Human Blood and Oral Fluid. Journal of Analytical Toxicology 2019; 43(4): 233–258. doi:10.1093/jat/bky104.

27. Newmeyer MN, Swortwood MJ, Barnes AJ, et al. Free and Glucuronide Whole Blood Cannabinoids’ Pharmacokinetics after Controlled Smoked, Vaporized, and Oral Cannabis Administration in Frequent and Occasional Cannabis Users: Identification of Recent Cannabis Intake. Clinical Chemistry 2016; 62(12): 1579–1592. doi:10.1373/clinchem.2016.263475.

28. Baglot SL, Hume C, Petrie GN, et al. Pharmacokinetics and central accumulation of delta-9-tetrahydrocannabinol (THC) and its bioactive metabolites are influenced by route of administration and sex in rats. Sci Rep 2021; 11(1): 23990. doi:10.1038/s41598-021-03242-7.

29. Torrens A, Roy P, Lin L, et al. Comparative Pharmacokinetics of Δ(9)-Tetrahydrocannabinol in Adolescent and Adult Male and Female Rats. Cannabis Cannabinoid Res 2022. doi:10.1089/can.2021.0205.

30. Nguyen JD, Aarde SM, Vandewater SA, et al. Inhaled delivery of Delta(9)-tetrahydrocannabinol (THC) to rats by e-cigarette vapor technology. Neuropharmacology 2016; 109: 112–120. doi:10.1016/j.neuropharm.2016.05.021.

31. Ruiz CM, Torrens A, Lallai V, et al. Pharmacokinetic and pharmacodynamic properties of aerosolized (“vaped”) THC in adolescent male and female rats. Psychopharmacology (Berl) 2021; 238(12): 3595–3605. doi:10.1007/s00213-021-05976-8.

32. Freels TG, Baxter-Potter LN, Lugo JM, et al. Vaporized Cannabis Extracts Have Reinforcing Properties and Support Conditioned Drug-Seeking Behavior in Rats. Journal of Neuroscience 2020; 40(9): 1897–1908. doi:10.1523/JNEUROSCI.2416-19.2020.

33. Glodosky NC, Cuttler C, Freels TG, et al. Cannabis vapor self-administration elicits sex- and dose-specific alterations in stress reactivity in rats. Neurobiol Stress 2020; 13: 100260. doi:10.1016/j.ynstr.2020.100260.

34. Hempel BJ, Wakeford AG, Nelson KH, et al. An assessment of sex differences in Delta(9)-tetrahydrocannabinol (THC) taste and place conditioning. Pharmacology Biochemistry and Behavior 2017; 153: 69–75. doi:10.1016/j.pbb.2016.11.006.

35. Cooper ZD and Craft RM. Sex-Dependent Effects of Cannabis and Cannabinoids: A Translational Perspective. Neuropsychopharmacology 2018; 43(1): 34–51. doi:10.1038/npp.2017.140.

36. Fattore L, Spano MS, Altea S, et al. Cannabinoid self-administration in rats: sex differences and the influence of ovarian function. British Journal of Pharmacology 2007; 152(5): 795–804. doi:10.1038/sj.bjp.0707465.

37. Fattore L, Spano MS, Altea S, et al. Drug- and cue-induced reinstatement of cannabinoid-seeking behaviour in male and female rats: influence of ovarian hormones. British Journal of Pharmacology 2010; 160(3): 724–35. doi:10.1111/j.1476-5381.2010.00734.x.

38. Wiley JL, Lefever TW, Marusich JA, et al. Comparison of the discriminative stimulus and response rate effects of (Delta9)-tetrahydrocannabinol and synthetic cannabinoids in female and male rats. Drug and Alcohol Dependence 2017; 172: 51–59. doi:10.1016/j.drugalcdep.2016.11.035.

39. Valjent E and Maldonado R. A behavioural model to reveal place preference to delta 9-tetrahydrocannabinol in mice. Psychopharmacology (Berl) 2000; 147(4): 436–8. doi:10.1007/s002130050013.

40. Ravula A, Chandasana H, Jagnarine D, et al. Pharmacokinetic and Pharmacodynamic Characterization of Tetrahydrocannabinol-Induced Cannabinoid Dependence After Chronic Passive Cannabis Smoke Exposure in Rats. Cannabis Cannabinoid Res 2019; 4(4): 240–254. doi:10.1089/can.2019.0049.

41. Sholler DJ, Strickland JC, Spindle TR, et al. Sex differences in the acute effects of oral and vaporized cannabis among healthy adults. Addiction Biology 2021; 26(4): e12968. doi:10.1111/adb.12968.

42. Macuchova E, Sevcikova M, Hrebickova I, et al. How various drugs affect anxiety-related behavior in male and female rats prenatally exposed to methamphetamine. International Journal of Developmental Neuroscience 2016; 51: 1–11. doi:10.1016/j.ijdevneu.2016.04.001.

43. Kuzmin A, Sandin J, Terenius L, et al. Acquisition, expression, and reinstatement of ethanol-induced conditioned place preference in mice: effects of opioid receptor-like 1 receptor agonists and naloxone. Journal of Pharmacology and Experimental Therapeutics 2003; 304(1): 310–8. doi:10.1124/jpet.102.041350.

44. Li X, Meng L, Huang K, et al. Environmental enrichment blocks reinstatement of ethanol-induced conditioned place preference in mice. Neuroscience Letters 2015; 599: 92–6. doi:10.1016/j.neulet.2015.05.035.

45. Cunningham CL, Ferree NK, and Howard MA. Apparatus bias and place conditioning with ethanol in mice. Psychopharmacology (Berl) 2003; 170(4): 409–22. doi:10.1007/s00213-003-1559-y.

46. Roma PG and Riley AL. Apparatus bias and the use of light and texture in place conditioning. Pharmacology Biochemistry and Behavior 2005; 82(1): 163–9. doi:10.1016/j.pbb.2005.08.004.

47. Calcagnetti DJ and Schechter MD. Extinction of cocaine-induced place approach in rats: a validation of the “biased” conditioning procedure. Brain Research Bulletin 1993; 30(5-6): 695-700. doi:10.1016/0361-9230(93)90102-h.

