## Supplemental Info for "Δ^9^-tetrahydrocannabinol (THC) vapor exposure produces conditioned place preference in male and female rats"

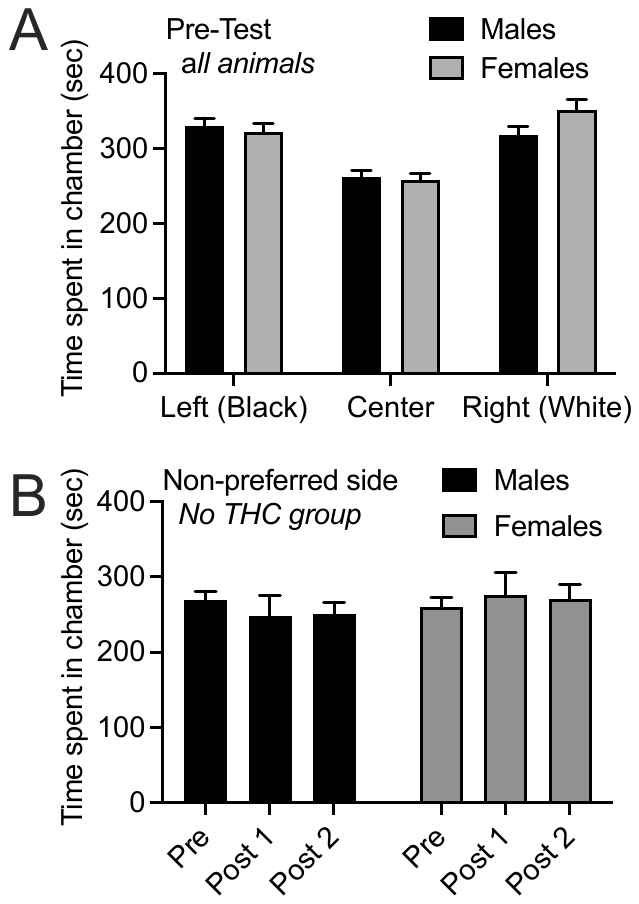

Supplemental Figure S1. Time spent in the 3 chambers in the Pre-test (A). Pre- and post-conditioning time spent in the non-preferred chambers in Vehicle vapor (no THC) exposed male and female rats (B).

| **Post Test 2** | Time Spent (s) | | | | Entries | | |
| --- | --- | --- | --- | --- | --- | --- | --- |
| Sex | Group | Preferred/ Veh-Paired | Center | Non-preferred/ Drug-Paired | Preferred/ Veh-Paired | Center | Non-preferred/ Drug-Paired |
| Males | No THC | 345.2 (16.8) | 303.9 (18.5) | 250.9 (15.4)^†^ | 64.4 (3.8) | 103.9 (7.3) | 61.8 (4.0) |
|  | Low THC | 389.4 (20.6) | 245.0 (15.1) | 265.6 (14.6)^†^ | 61.8 (4.5) | 95.9 (5.1) | 55.9 (2.5) |
|  | Med THC | 310.7 (22.9) | 268.5 (22.7) | 320.8 (24.7) | 65.8 (4.4) | 102.5 (5.7) | 59.1 (3.4) |
|  | High THC | 282.0 (16.6) | 312.9 (18.6) | 305.0 (9.0) | 61.3 (2.2) | 100.3 (3.0) | 61.8 (2.4) |
| Females | No THC | 364.8 (36.9) | 264.2 (24.5) | 271.0 (19.3)^†^ | 53.6 (3.7) | 77.1 (5.4) | 48.0 (2.2) |
|  | Low THC | 355.0 (14.2) | 302.8 (16.6) | 242.2 (13.5)^†^ | 59.1 (1.9) | 94.3 (3.7) | 51.3 (3.5) |
|  | Med THC | 404.0 (25.5) | 243.7 (16.8) | 252.4 (13.4)^†^ | 60.1 (3.1) | 85.3 (4.7) | 52.4 (2.0) |
|  | High THC | 288.4 (26.3) | 285.8 (16.7) | 325.8 (15.6) | 57.8 (3.4) | 100.0 (5.2) | 64.0 (4.8) |

Supplemental Table S1. Time spent (in seconds; mean (SEM)) and number of entries in each chamber during post-test 2. † indicates less time spent in that chamber compared with the initially preferred chamber.
